# Rule adherence warps decision-making

**DOI:** 10.1101/2019.12.16.878306

**Authors:** R. Becket Ebitz, Jiaxin Cindy Tu, Benjamin Y. Hayden

**Author notes:** Corresponding author: Becket Ebitz, Department of Neuroscience, University of Minnesota, Minneapolis MN 55455.

## Abstract

We have the capacity to follow arbitrary stimulus-response rules, meaning policies that determine how we will behave across circumstances. Yet, it is not clear how rules guide sensorimotor decision-making in the brain. Here, we recorded from neurons in three regions linked to decision-making, the orbitofrontal cortex, ventral striatum, and dorsal striatum, while macaques performed a rule-based decision-making task. We found that different rules warped the neural representations of chosen options by expanding rule-relevant coding dimensions relative to rule-irrelevant ones. Some cognitive theories suggest that warping could increase processing efficiency by facilitating rule-relevant computations at the expense of irrelevant ones. To test this idea, we modeled rules as the latent causes of decisions and identified a set of “rule-free” choices that could not be explained by simple rules. Contrasting these with rule-based choices revealed that following rules decreased the energetic cost of decision-making while warping the representational geometry of choice.

**SIGNIFICANCE STATEMENT:** One important part of our ability to adapt flexibly to the world around us is our ability to implement arbitrary stimulus-response mappings, known as “rules”. Many studies have shown that when we follow a rule, its identity is encoded in neuronal firing rates. However, it remains unclear how rules regulate behavior. Here, we report that rules warp the way that sensorimotor information is represented in decision-making circuits: enhancing information that is relevant to the current rule at the expense of information that is irrelevant. These results imply that rules are implemented as a kind of attentional gate on what information is available for decision-making.

## INTRODUCTION

We have a tremendous capacity to change how we perceive and respond to the world as our environment and goals change. One central part of this flexibility is our ability to implement stimulus-response rules: policies for guiding behavior that allow us to make decisions quickly, reasonably accurately, and with little subjective sense of effort (Asaad et al., 2000; Bunge, 2004; Wallis et al., 2001; White & Wise, 1999). We know from many studies that rule identity is encoded in firing rate changes of neurons in specific brain regions, classically in the dorsolateral prefrontal cortex (dlPFC, Asaad et al., 2000; Bunge, 2004; Wallis et al., 2001; White & Wise, 1999), but also in many other regions implicated in decision-making, like the orbitofrontal cortex (OFC) and striatum (Bissonette & Roesch, 2015; Sleezer et al., 2016; Sleezer et al., 2017; Sleezer & Hayden, 2016; Tsujimoto et al., 2011; Yamada et al., 2010). However, implementing a rule requires more than simply encoding its identity in the right structures and an important question remains unanswered: how do rules shape neural processing in order to control behavior?

We hypothesized that the brain may implement rules, in part, as a kind of attentional gate that warps the way that information is represented in sensorimotor neurons. In this view, rules depend on a top-down biasing signal that enhances the representation of relevant information at the expense of irrelevant information, warping the geometry of our representations of the world. This basic model is at the heart of influential views of not just rule-guided behavior, but of executive function in general (Miller & Cohen, 2001). However, tests of this hypothesis in both dorsolateral prefrontal cortex (Lauwereyns et al., 2001; Mante et al., 2013) and early sensory cortex (Katzner et al., 2009; Sasaki & Uka, 2009) have failed to find evidence that rules are implemented through attentional gating. Of course, while negative results like these are suggestive, they cannot rule out the possibility that rules do gate information in other circuits.

Within decision-making circuits, the idea that rules may warp representations is particularly compelling because this warping could, in theory, streamline decision-making. Many decisions we make are not based on simple rules. Instead, decision-making often requires us to solve very high-dimensional problems, where the correct response is a complex function of our past choices, reward history, and the options available to us in the moment (Simon, 1955). However, an attentional gate could substantially reduce the effective dimensionality of decision-making problems by reshaping the way we represent the world to highlight only a subset of the dimensions of the problem: the rule-relevant information (Gigerenzer & Todd, 1999; Shah & Oppenheimer, 2008; Tversky, 1972). Thus, if rules did gate how choice information is represented in the brain and in decision-making circuits, then rule-based decisions could be a more efficient use of limited neural resources than decisions that do not rely on a rule (Gigerenzer & Gaissmaier, 2011; Gigerenzer & Todd, 1999; Sanfey et al., 2006; Shah & Oppenheimer, 2008; Simon, 1955). However, while this idea makes intuitive sense and may align to our introspective experience, it has not been tested empirically.

Of course, rule-based decisions could be more efficient than rule-free decisions for reasons other than representational warping. An alternative idea is that rule-based decisions depend on a functional handoff from computationally inefficient, deliberative decision-making structures to more efficient, automatic ones (Balleine & O’Doherty, 2010; Frank, Cohen, & Sanfey, 2009; Sanfey et al., 2006; van der Meer et al., 2012). We call this view the ***handoff hypothesis***, to differentiate it from our ***warping hypothesis***. This view is based on observations that some structures, like the dorsal striatum (DS), are implicated in rapid stimulus-bound automatic decisions (Balleine et al., 2007; Jog et al, 1999; Yin & Knowlton, 2006), while other structures, like the ventral striatum (VS) and the orbitofrontal cortex (OFC), are implicated in more flexible, deliberative decision-making (Buckley et al., 2009; O’Doherty et al., 2004; Rudebeck & Murray, 2014; Valentin et al., 2007; Walton et al., 2010). However, this hypothesis also remains untested, in large part because of the difficulty of identifying when a decision is, or is not, rule-based.

Here, we used a novel mathematical technique to infer the hidden rules underlying decisions and determine 1) if rule-based decisions are an efficient use of limited neural resources, and, 2) if so, why. To do this, we recorded from Area 13 of the OFC, the nucleus accumbens core of the VS, and both the caudate and putamen in the DS in a macaque analog of the Wisconsin Card Sort Task. Previous studies have generated insight into the neural basis of rules through comparing periods of correct task performance in which one rule or another was imposed by the task (Asaad et al., 2000; Bunge, 2004; Everling & DeSouza, 2005; Mante et al., 2013; Siegel et al., 2015; Sleezer et al., 2016; Wallis et al., 2001; White & Wise, 1999; Womelsdorf et al., 2010). However, here, we compare periods in which monkeys were following one rule or another, regardless of whether that was correct. We do this by modeling rules as the latent stimulus-response policies underlying the monkeys’ decisions (Ebitz et al., 2019).

## RESULTS

Two rhesus macaques performed a total of 128 sessions (73,627 trials) of the Cognitive Set-Shifting Task (CSST, **Figure 1A**, (Moore et al., 2005; Sleezer et al., 2016), a macaque analog of the Wisconsin Card Sorting task that encourages subjects to discover and apply hidden stimulus-response rules to make decisions. On each trial, a unique combination of the three colored (cyan, magenta, yellow) shapes (circle, star, triangle) appeared in random order at three screen locations. On each trial, there was a correct color or shape (six possible “correct features”). Only choices that matched the correct feature were rewarded. The correct feature was fixed for a block of 15 correct trials, then it changed and a new correct feature was chosen at random (See **Methods**). When the subjects discovered the correct feature, they had the opportunity to make rule-based decisions by following a rule where they only chose options that matched the correct feature, generating a sequence of choices that shared a single color or shape feature (**Figure 1B**). However, after the correct feature changed, subjects had to discard any rule they had been following and focus on discovering the new correct rule.

**Figure. 1.**
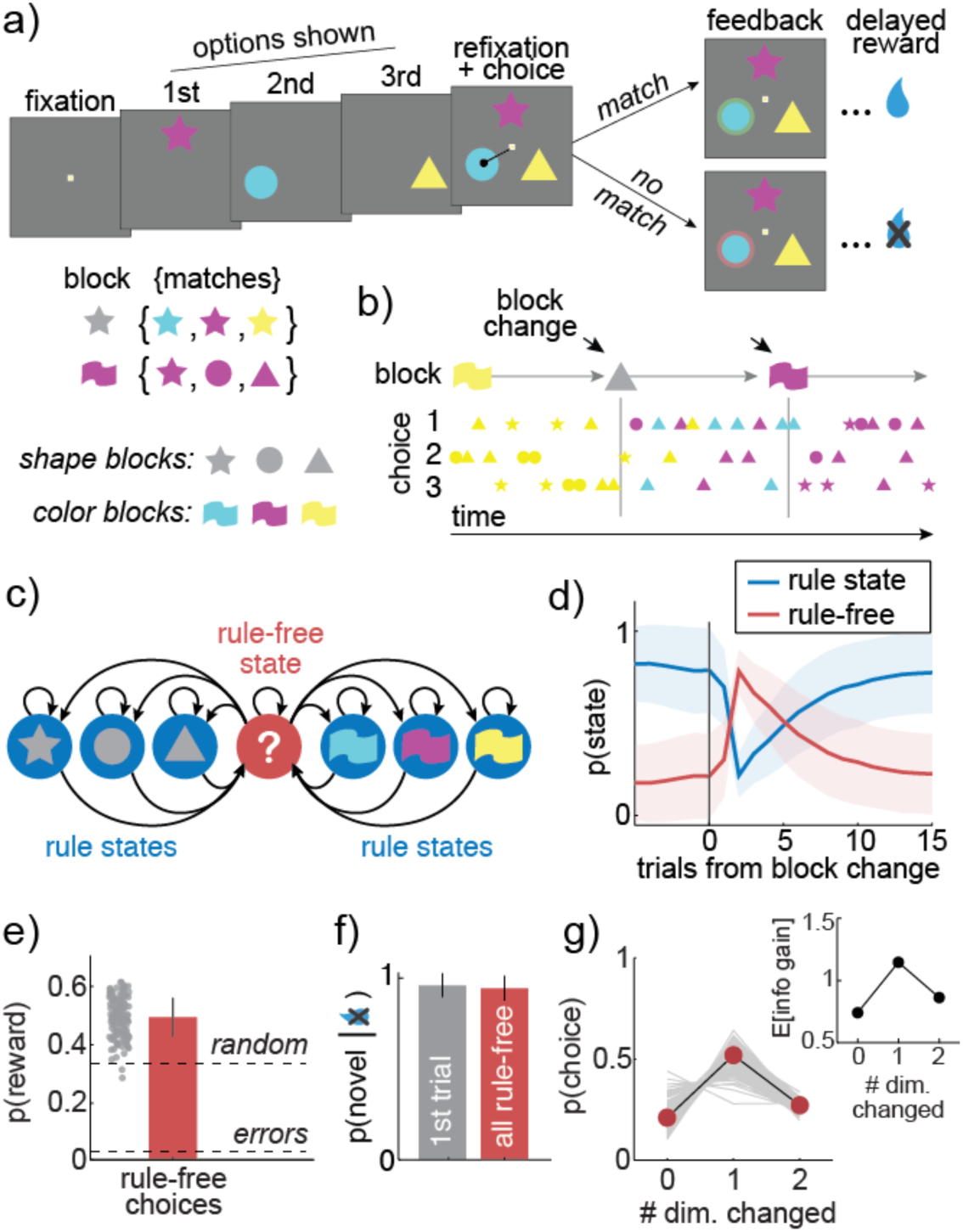
Task design and latent state identification. A) Trial structure in the cognitive set shifting task. Three options are presented sequentially and the subject indicates choice by refixating a central fixation point, then shifting gaze to one option. Visual feedback is displayed and then the subject is rewarded if he chose an option that matched the current block. Inset) There are six possible blocks, three shape blocks (star, circle, triangle) and three color blocks (cyan, magenta, yellow). Correct choices match the block’s rewarded feature, but differ in the irrelevant feature domain. B) An example sequence of trials, illustrating the blocks (top row). Each symbol reflects the color and shape of one choice. C) The hidden Markov model used to identify the latent states underlying decisions. Each rule-based state produced choices that matched the state’s feature. Conversely, the rule-free state produced all choices uniformly. D) The mean probability of rule-based and rule-free states, aligned to the block changes. Shading = STD across sessions. E) Accuracy of rule-free choices on average (bar) and within individual sessions (gray dots). Accuracy was greater than expected from errors (bottom dashed line) or random choice (top dashed line). Bars = STD across sessions. F) Probability of choosing a novel option (matching neither the last choice’s shape or color) after the first omitted reward after a block change (gray) and for all rule-free choices that followed no-reward outcomes (red). Bars = STD across sessions. G) The probability of choosing options that differ in 0, 1, or 2 feature dimensions from the last choice on average (red markers) and within individual sessions (gray lines). Compare with the expected information gain about what rule is best for each choice types (inset), calculated from the history of choices and rewards observed in rule-free trials.

To infer what rule, if any, subjects were applying on each trial, we used a hidden Markov model (HMM; see **Methods**; **Figure 1C**). The HMM modelled each rule as a latent state that produced choices that matched the rule’s relevant feature (i.e. the blue rule produced blue choices, regardless of shape). The model allowed us to make statistical inferences about what specific rule was underlying each choice (i.e. to determine if a blue-star choice was due to a blue rule or a star rule). The rules inferred by the HMM matched the current correct feature 93% of the time on average (± 2% STD, 95% CI across sessions: 89.5% to 97.4%). Thus, the model identified the periods when animals were following rules that very often (though not always) matched the rule imposed by the task. In addition to 6 rule states, the model also included a “rule-free” state to account for the fact that some choices could not be explained by a simple sensorimotor rule (Ebitz et al., 2019). Rule-free choices occurred most frequently, but not exclusively, at block changes (**Figure 1D**), indicating that the non-rule-based choices tended to occur precisely when the subjects did not know what rule to follow. However, non-rule-based choices also occurred at times when monkeys could have been following rules but were not. For example, 19.5% of the correct choices that occurred during stable periods (>5 trials after block changes) were not rule-based choices per the HMM (± 18% STD, 95% CI across sessions: 1% to 62%). Overall, 67% of trials were classified as rule-based (n= 49,043/73,627) while 33% were classified as rule-free (n= 24,584/73,627).

In previous studies, decisions that do not match the current rule have been dismissed as errors. However, rule-free decisions were often correct choices here. In fact, the rule-free decisions identified by the HMM were strategic, deliberative decisions that maximized both reward and information about what rule was best. For example, rule free decisions were correct 49% of the time (± 6.9% STD), far more frequently than we would expect from either errors or random decisions (**Figure 1E**). In comparison, errors would be correct approximately 3% of the time (the probability that an error would coincide with a block change and target the new correct feature; significant difference: p <0.0001, t(1,127) = 77, one-sample t-test) and random decisions would only be correct 33% of the time (p < 0.0001, t(1,127) = 26.9, one-sample t-test). This is impressive because rule-free choices occurred most frequently after block changes, when the animals could not know what the correct feature was.

Rule-free decisions tended to be correct because these decisions were made strategically. For example, after an unexpected reward omission, choosing a novel option—one that differs in both color and shape from the last choice—maximizes both reward likelihood and information about the correct feature. On the first trial after feedback that the correct feature has changed, subjects made this optimal choice 95% of the time (± 6.7% STD across sessions, sig. different than chance, p < 0.0001, t(1,127) = 50.6). Similarly, across all rule-free trials where the subjects were not rewarded on the last trial, they made the same novel choice 94% of the time (± 6.8%; **Figure 1F**; p < 0.0001, t(1,127) = 46.7). Moreover, rule-free choices in general were strongly biased towards the options that would maximize information, as calculated from the the actual pattern of choices and rewards that preceded these choices (**Figure 1G**; see Methods). Thus, rule-free choices used a strategy that integrated information about past rewards and choices to maximize both reward and information about what rule was best.

### Rule adherence reduces firing rate

We recorded responses of individual neurons in OFC (n = 115 cells), VS (n = 103), and DS (n = 204; recording sites in **Figure 2A**). We first looked for hypothesized differences in overall spiking activity during rule-based and rule-free decisions. Across the option viewing and choice period (before feedback), we found that firing rates were systematically lower during rule-based decisions in all three regions (example cells: **Figure 2B**; population: **Figure 2C**; VS: reduction of 0.38 spikes/sec ± 0.14 STE across neurons, p < 0.0001; DS, reduction of 0.19 ± 0.10 spikes/sec, p < 0.0001; OFC, reduction of 0.17 ± 0.11 spikes/sec, p < 0.0001; permutation test against expected difference with shuffled state labels). This was not due to differences in reward expectation between the two conditions because reward-dependent changes in firing rate were both smaller and in the opposite direction in all three regions (no reward - reward, VS: −0.28 ± 0.09, DS: −0.09 ± 0.07, OFC: −0.06 ± 0.07). Thus, if anything, we would be underestimating the extent to which adhering to a rule decreases spiking activity in these regions because rule-based trials were more likely to be rewarded than rule-free trials.

**Figure. 2.**
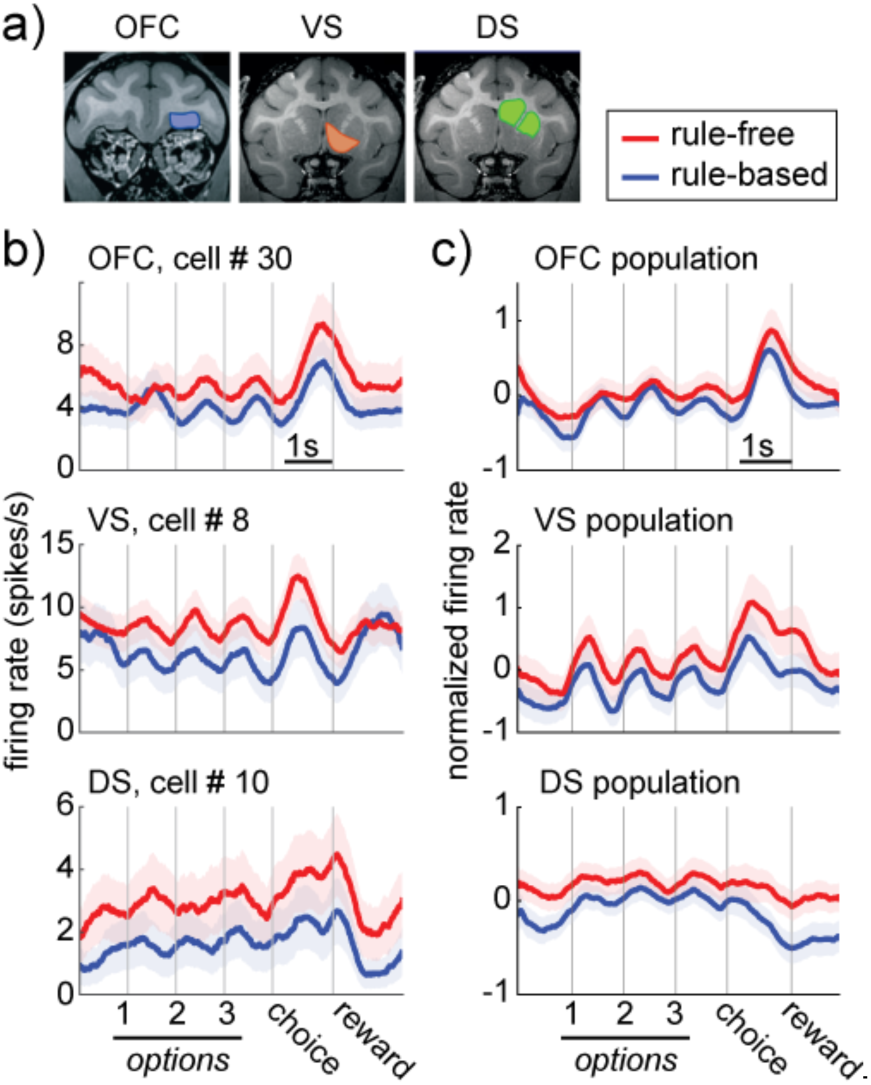
Firing rate decreases during rule-based decisions. A) Location of recording sites in the OFC, VS, and DS. B) Example cells in the OFC (top), VS (middle), and DS (bottom), recorded during rule-free (red) and rule-based decisions (blue). Bars = standard errors across trials. C) Same as B for the populations. Bars = standard errors across neurons.

### Rule adherence increases information about choice identity

Neurons in all three areas encoded the visual features of the upcoming choice (“choice identity”; **Figure 3A and B**; one-way ANOVAs, predicting choice identity from the firing rate of each neuron; VS: n = 52/103, proportion = 0.50, p <0.0001; DS: n = 85/204; proportion = 0.41, p <0.0001; OFC: n = 52/115, proportion = 0.45, p < 0.0001; one-sided binomial test for difference from expected false positive rate of 0.05). If rule-based decisions are more efficient than rule-free because of an attentional gate that warps the information available for decision-making, then we would see distributed changes in decision-making efficiency across all regions. The rate of choice-predictive information per spike should increase during rule use across all regions (**Figure 3C, top**). Conversely, if rule-based decisions are more efficient due to a handoff from deliberative to automatic decision-making structures (**Figure 3C, bottom**), then choice-predictive information should selectively decrease in regions implicated in deliberative decision-making, as these regions are less involved in decision-making.

**Figure 3.**
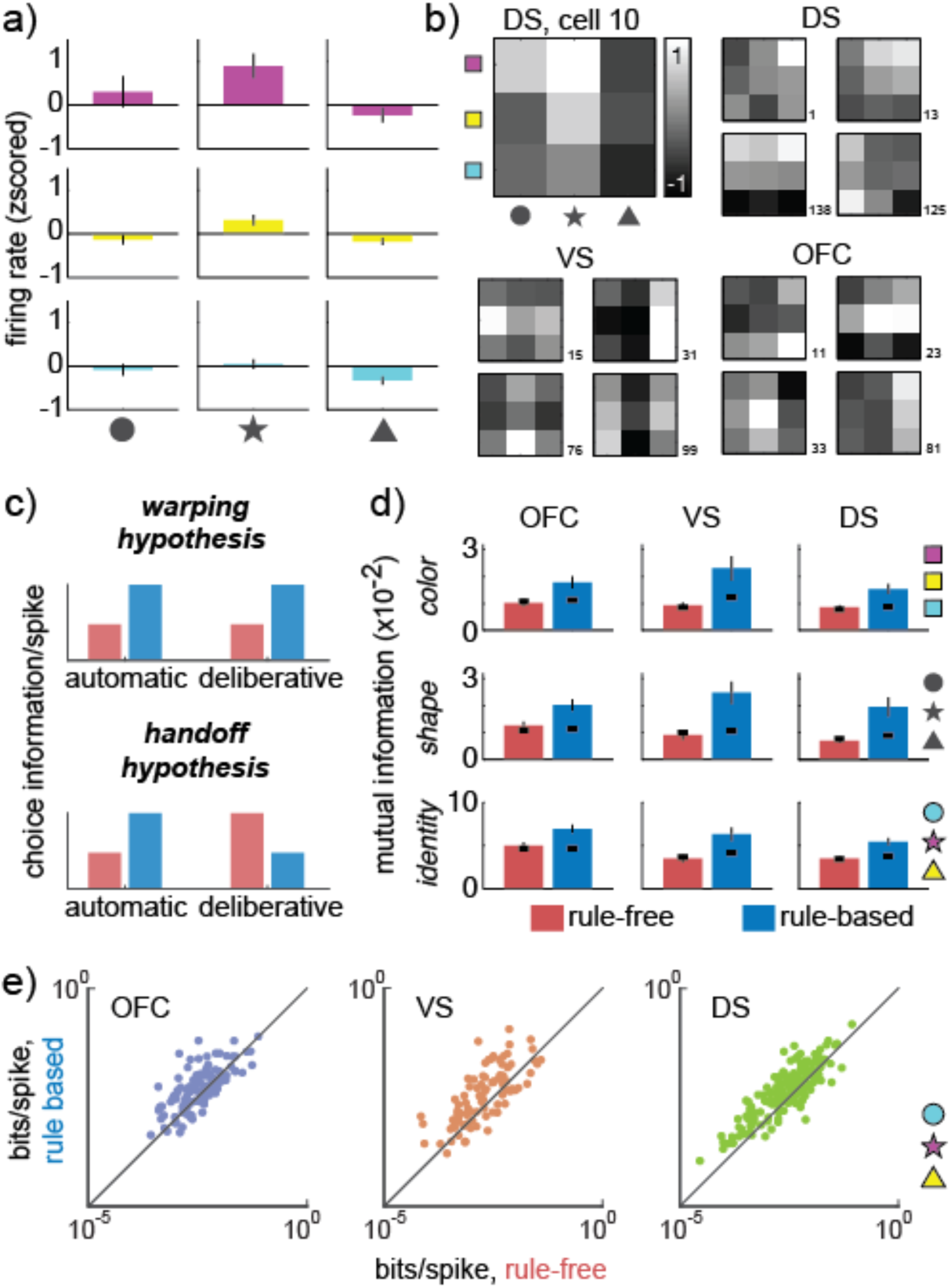
Choice information increases during rule-based decisions. A) Choice identity tuning of an example neuron recorded from DS. B) The same example neuron represented as a choice tuning map (top left), and the tuning maps of four additional example neurons from DS, VS, and OFC. C) The two hypotheses make different predictions for how choice information per spike (efficiency) should change as a function of whether neurons are in automatic (left) or deliberative (right) decision-making regions. D) Information about the color (top row), shape (middle row), and identity (bottom row) of the chosen option in each of OFC (left column), VS (middle column), and DS (right column). Bars = STE across neurons and thick black lines = shuffled data. E) Choice identity information per spike during rule-based (y-axis) and rule-free decisions (x-axis), measured in individual cells (dots) recorded in each of OFC (left, purple), VS (middle, orange), and DS (right, green). Dots on the unity line indicate no change in bits/spike across rule-free and rule-based decisions, dots above unity indicate an increase in bits/spike during rule-based decisions. Log scale.

To compare the amount of choice-predictive information across rule-based and rule-free decisions, we calculated the mutual information between each neuron’s firing rate and the chosen option separately for each condition (see **Methods**). Choice-predictive information was increased during rule-based decisions, compared to rule-free decisions, in VS (rule-based: average of 0.06 bits per trial ± 0.008 SEM, rule-free: 0.03 ± 0.004, p < 0.0001, z = 4.40, n = 103, Wilcoxon rank sum test), DS (rule-based: 0.05 ± 0.004, rule-free: 0.03 ± 0.003, p < 0.0001, z = 6.21, n = 204), and OFC (rule-based: 0.07 ± 0.005, rule free: 0.05 ± 0.004, p < 0.0002, z = 3.77, n = 115). Although mutual information was lower during rule-free decisions, some neurons did still encode choice identity (one-way ANOVAs, predicting chosen stimulus identity from the firing rate of each neuron; VS: n = 9/103, proportion = 0.09, p <0.04; DS: n = 23/204, proportion = 0.11, p <0.0001; OFC: n = 14/115, proportion = 0.12, p < 0.001; one-sided binomial test for difference from expected false positive rate of 0.05). Thus, choice information was increased during rule-based decisions, but it was not completely absent during rule-free decisions.

The increase in choice information during rule-based decisions was not a trivial consequence of the difference in firing rate between the two types of decisions. First, mutual information typically *increases* with mean firing rate (Panzeri et al., 2007), but firing rates were lower during rule-based decisions than rule-free ones. Thus, if anything, we are likely underestimating the magnitude of the increase in choice information during rule-based decisions. Second, the total information per spike also increased during rule-based decisions in all three regions (VS, rule-based: 0.012 bits per spike ± 0.005, rule-free: 0.004 ± 0.001, p < 0.0002, z = 3.84, DS, rule-based: 0.008 ± 0.001, rule-free: 0.005 ± 0.0006, p < 0.0001, z = 6.11, and OFC, rule-based: 0.009 ± 0.001, rule-free: 0.007 ± 0.001, p < 0.002, z = 3.11). Thus, computational efficiency increased during rule-based decisions in all three regions.

### Rule adherence has different effects on sparsity across regions

The distributed firing rate effects we observed differ qualitatively from the results of fMRI studies, which tend to support the idea of a functional handoff because changes in the BOLD signal are observed across different regions during different types of decision-making (O’Doherty et al., 2004; Poldrack et al., 2001; Valentin et al., 2007). We hypothesized that the apparent discrepancy between these results could be due to fact that the BOLD signal may reflect population measures other than average spiking, such as how densely or sparsely activity is distributed across a population of neurons (Logothetis, 2008).

Therefore, we next measured the sparsity of activity across VS, DS, and OFC during rule-based and rule-free decision-making with the Gini index (**Figure S1B**; see **Methods**; Hurley & Rickard, 2009). In VS, the population response was more sparse during rule-based decisions (Gini index = 0.662) than rule-free decisions (0.656, two-sample t-test, p < 0.004, t(1,358) = 2.92, 95% CI for the difference = [0.002, 0.009]). However, there was an opposite pattern in both DS (rule-based = 0.690, rule-free = 0.696, p < 0.0001, t(1,358) = −4.69, 95% CI = [−0.009, − 0.004]) and OFC (rule-based = 0.541, rule-free = 0.562, p < 0.0001, t(1,358) = −9.54, 95% CI = [−0.03, −0.02]). Thus, while spiking was lower during rule-based decisions in all three regions, spikes were more densely distributed in VS during rule-free decision-making, but more densely distributed in OFC and DS during rule-based decision-making.

### Rule adherence warps choice representations

Together, these results broadly support the warping hypothesis, rather than the handoff hypothesis. However, the warping hypothesis predicts that rule-based decisions should be more efficient than other decisions because of changes in how choices are represented in decision circuits. Though we cannot directly ask how different goal states change cognitive representations (Decharms & Zador, 2000), we can take advantage of the fact that neural representations have metric relationships to each other that recapitulate cognitive similarities (Kriegeskorte et al., 2008; Shepard & Chipman, 1970). This means that we can ask how the similarity of neural representations of different choices changes under different goal states (see **Methods**, (Cukur, Nishimoto, Huth, & Gallant, 2013)). Here, we define a choice representation within decision-making regions as a pattern of firing rates across a population of neurons, or equivalently, as vector in neuron-dimensional state space (**Figure 4A**). Because neurons were recorded largely separately, we built representational vectors from pseudopopulations (see **Methods**; (Churchland et al., 2012; Machens et al., 2010; Mante et al., 2013; Meyers et al., 2008).

**Figure 4:**
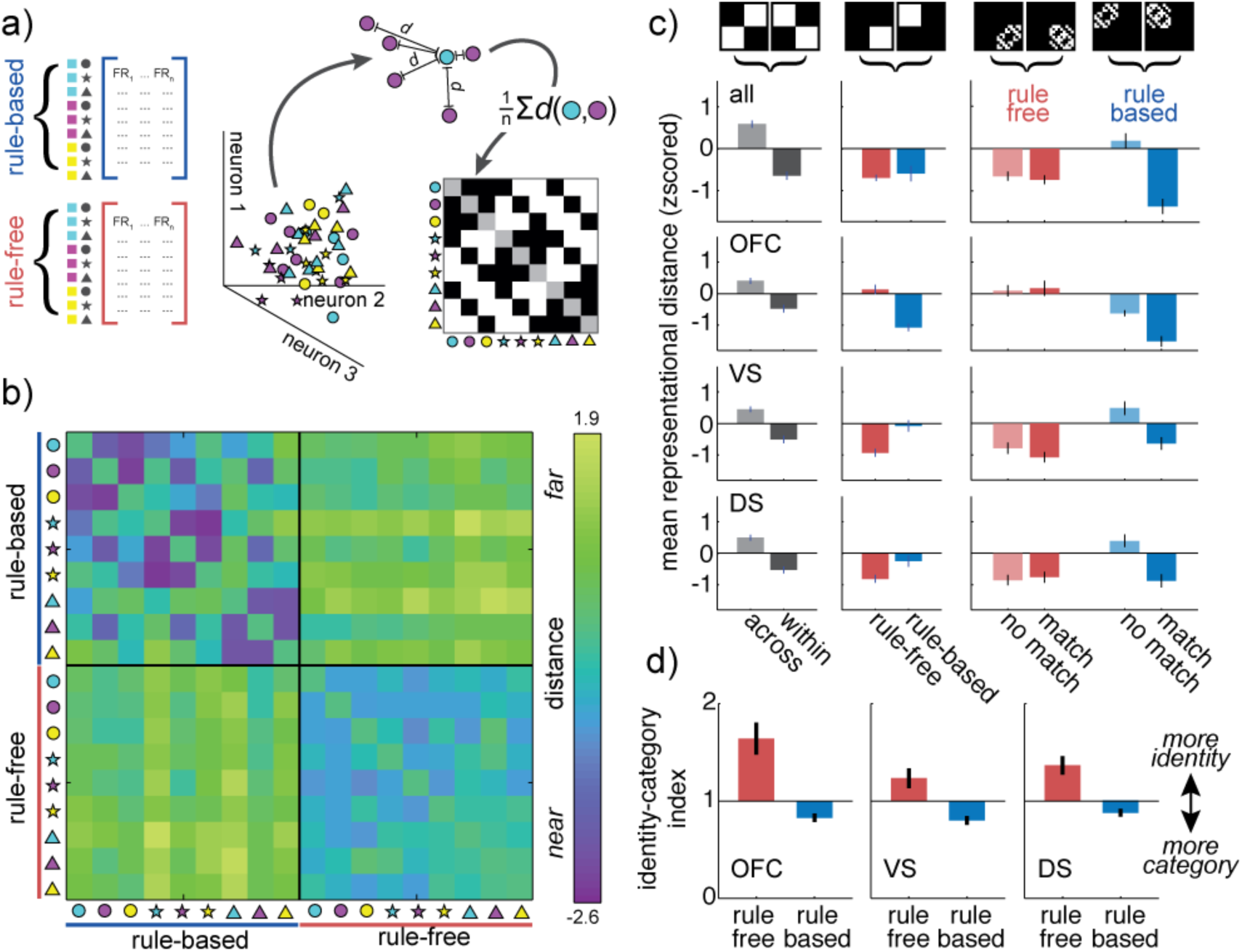
Changes in choice representations across rule-based and rule-free decision-making. A) Separately recorded neurons are combined into rule-based and rule-free pseudopopulations (left) in order to estimate the distribution of choice representation “locations” in neuron-dimensional space (middle). The average distance between different types of choices (here, between cyan circle and magenta circles) is calculated and used to populate a representational similarity matrix (right). The example matrix is the expected structure if choices that share a color or shape are closer together in neuron-dimensional space than choices that do not share features. B) The representational similarity matrix for both rule-based (upper left) and rule-free (bottom-right) decisions, calculated across all regions. C) Contrasts across the representational similarity matrix. Left column) Any change in representation between rule-based and rule-free decisions would increase distance between representations measured across decision types (light gray), compared to within decision types (dark gray). Middle column) A change in the total representational space between decision types would appear as a difference in the mean distance within rule-based decisions (blue) compared to rule-free decisions (red). Right column) Mean distance between stimuli that share features (“match”, dark color) and do not share features (“no match”, light color), for both rule-free (red) and rule-based (blue) decisions. A decrease in the match distance, relative to no-match, indicates an increase in categorical structure. Bars = STE across unique matrix elements. D) Same as C, separately for OFC (top row), VS (middle) and DS (bottom). E) The identity-category index for single neurons in OFC (left), VS (middle) and DS (left). Bars = STE across neurons.

First, we asked whether there were representational changes due to rule-based decision-making in general, rather than changes due to applying a specific rule. Because we had no knowledge of which, if any, stimulus feature was focal during rule-free decisions, the rule-free pseudopopulation necessarily combined information across trials in which different stimulus features were focal. To create an equivalent rule-based pseudopopulation, rule-based trials were sampled without regard for the rule. We first combined the data across VS, DS, and OFC, asking how the distances between distributed choice representations changed across choices and goals (Cukur et al., 2013). We illustrate the pairwise distance between the representations of all possible choices during both rule-based and rule-free decision-making as a representational similarity matrix (**Figure 4B**; see Methods).

There was a representational shift between rule-based and rule-free decisions such that similar choices were represented by more similar patterns of activity during rule-based decisions. This representational shift was due to three changes in the representational similarity matrix (**Figure 4C**). First, choice representations changed across rule state. Choice representations were closer to themselves within choice type than between choice types (within: mean distance = − 0.64 ± 0.10 STE, between = 0.59 ± 0.09, p < 0.0001, t(1,71) = 9.62, 95% CI = [0.98, 1.49]). However, second, this was not due to a simple expansion of the representational space during rule-based decisions because the average distance between choice representations was unchanged across choice types (rule-based: mean distance = −0.59 ± 0.18, rule-free: mean distance = −0.69 ± 0.08, p > 0.5, t(1,35) = 0.57, 95% CI for effect size = [−0.26, 0.47], paired t-test). Third, choices that shared a feature were closer than choices that did not share a feature during rule-based decision-making (shared feature = −1.37 ± 0.18; no shared feature = 0.18 ± 0.18, p < 0.0001, t(1,34) = 6.12, 95% CI = [1.03, 2.06], two-sample t-test), but not during rule-free decision-making (shared feature = −0.74 ± 0.10; no shared feature = −0.65 ± 0.11, p > 0.5, t(1,34) = 0.58, two-sample t-test). Thus, choice representations shifted to become more organized with respect to stimulus categories during rule-based decisions.

We next asked whether these effects were the same across OFC, VS, or DS or driven by representational changes in one regions. Choices were more similar within rule-based or rule-free decisions than between choice types in all three regions, indicating that representational shifts did occur in each region (OFC: within = −0.47 ± 0.12 STE, between = 0.41 ± 0.10, p < 0.0001, t(1,71) = 7.16; VS: within = −0.50 ± 0.12, between = 0.45 ± 0.09, p < 0.0001, t(1,71) = 6.39; DS: within = −0.53 ± 0.11, between = 0.49 ± 0.09, p < 0.0001, t(1,71) = 7.74). However, the regions differed in how the total size of the representational space changed across choice types. In OFC, choice representations were closer together, on average, during rule-based decisions than rule-free decisions (rule-based = −1.07 ± 0.12, rule-free = 0.13 ± 0.15, p < 0.0001, t(1,35) = 6.48, paired t-test), but in VS and DS the representational space was expanded during rule-based decision-making, compared to rule-free (VS: rule-based = −0.08 ± 0.18, rule-free = − 0.93 ± 0.12, p < 0.0001, t(1,35) = 5.17; DS: rule-based = −0.25 ± 0.18, rule-free = −0.81 ± 0.12, p < 0.02, t(1,71) = 2.60). Nevertheless, in all three regions, choices that shared a feature were only closer to each other than choices that did not share a feature during rule-based decisions (shared feature = −1.52 ± 0.17, no shared feature = −0.62 ± 0.09, p < 0.0001, t(1,34) = 4.63, two-sample t-test; VS: shared feature = −0.64 ± 0.20, no shared feature = 0.48 ± 0.21, p < 0.001, t(1,34) = 3.82; DS: shared feature = −0.88 ± 0.21, no shared feature = 0.39 ± 0.20, p < 0.0002, t(1,34) = 4.40), but not during rule-free decisions (OFC: shared feature = 0.17 ± 0.24, no shared feature = 0.09 ± 0.18, p > 0.7, t(1,34) = 0.27, two-sample t-test; VS: shared feature = −1.07 ± 0.16, no shared feature = −0.79 ± 0.18, p > 0.2, t(1,34) = 1.19; DS: shared feature = −0.77 ± 0.18; no shared feature = −0.86 ± 0.16; p > 0.7, t(1,34) = 0.38). Thus, while choice representations were only more compact during rule-based decision-making in OFC—not in striatum—choice representations shifted so that they were more categorically organized during rule-based decision-making in all three regions.

### Single-neuron choice tuning emphasizes category, rather than identity, during rule-based decisions

Because the choice representations were constructed from pseudopopulations rather than simultaneously recorded populations, the increase in category encoding during rule-based decisions could have been an artifact of how we combined activity across neurons. However, representational changes at the population level should also be associated with changes in the tuning of single neurons (Cukur et al., 2013). Here, the changes in choice representations at the population level suggested that single neurons should have less tuning for choice identity (combination of color and shape), but more tuning for choice category (color or shape) during rule-based decisions.

We quantified the relative amount of choice identity and choice category tuning in single neurons with an “identity-category index” (see Methods). We found a significant decrease in the identity-category index (relative increase in categorization) during rule-based in all three regions (**Figure 4C**): in neurons recorded from OFC (rule-free = 1.64 ± 0.16 STE, rule-based = 0.83 ± 0.04, sig. decrease, p < 0.0001, t(1,114) = −4.77, paired t-test), VS (rule-free = 1.23 ± 0.10, rule-based = 0.80 ± 0.05, p < 0.0002, t(1,102) = −3.95) and DS (rule-free = 1.37 ± 0.09 STE, rule-based = 0.88 ± 0.04, p < 0.0001, t(1,202) = −4.59). Thus, during rule-based decisions, even single neurons shifted from encoding choice identity to encoding choice category.

### Color rules and shape rules have different effects on choice representations

Choice representations became more categorically organized during rule-based decision-making. This is consistent with the warping hypothesis because, if rule-irrelevant encoding dimensions were collapsed, choices that differed only in rule-irrelevant dimensions would be represented more similarly. To determine whether the categorical structure emerged from selective representational shifts along rule-relevant or irrelevant dimensions, we next performed similar analyses on pseudopopulations constructed separately for shape and color rules. Combining across OFC, VS, and DS, there was a representational shift between color-rule and shape-rule choice representations (mean distance within rule type = −0.33 ± 0.15 STE, between rule types = 0.28 ± 0.06; p < 0.002, t(1,71) = 3.41, 95% CI = [0.25, 0.96]). However, this representational shift specifically decreased the representational distance between rule-irrelevant features, both during the color rule (mean distance between same-color choices = 0.35 ± 0.22, same-shape = −2.43 ± 0.14, p < 0.0001, t(1,16) = 10.70, 95% CI = [2.24, 3.35]), and during the shape rule (mean distance between same-shape choices = 0.29 ± 0.09, same-color = −2.31 ± 0.14, p < 0.0001, t(1,16) = 15.62, 95% CI = [2.25, 2.96]). Thus, the categorical structure that emerged during rule-based decision-making was due to an increase in the representational similarity of choices that differed solely along rule-irrelevant dimensions, consistent with the predictions of the warping hypothesis.

We observed significant representational changes within each region individually: OFC (within rule types = −0.35 ± 0.14, between = 0.31 ± 0.08, p < 0.0002, t(1,71) = 4.01, 95% CI = [0.33, 0.99]), VS (within rule types = −0.21 ± 0.14, between = 0.19 ± 0.09, p < 0.03, t(1,71) = 2.27, 95% CI = [0.05, 0.75]), and DS (within rule types = −0.32 ± 0.15, between = 0.30 ± 0.06, p < 0.0005, t(1,71) = 3.72, 95% CI = [0.28, 0.96]). Moreover, within each region, the representational shifts were identical to what we observed when combining across regions (**Figure 5**; **Table S1**). Thus, these representational changes were broadly distributed across regions implicated in different types of decision-making.

**Figure 5:**
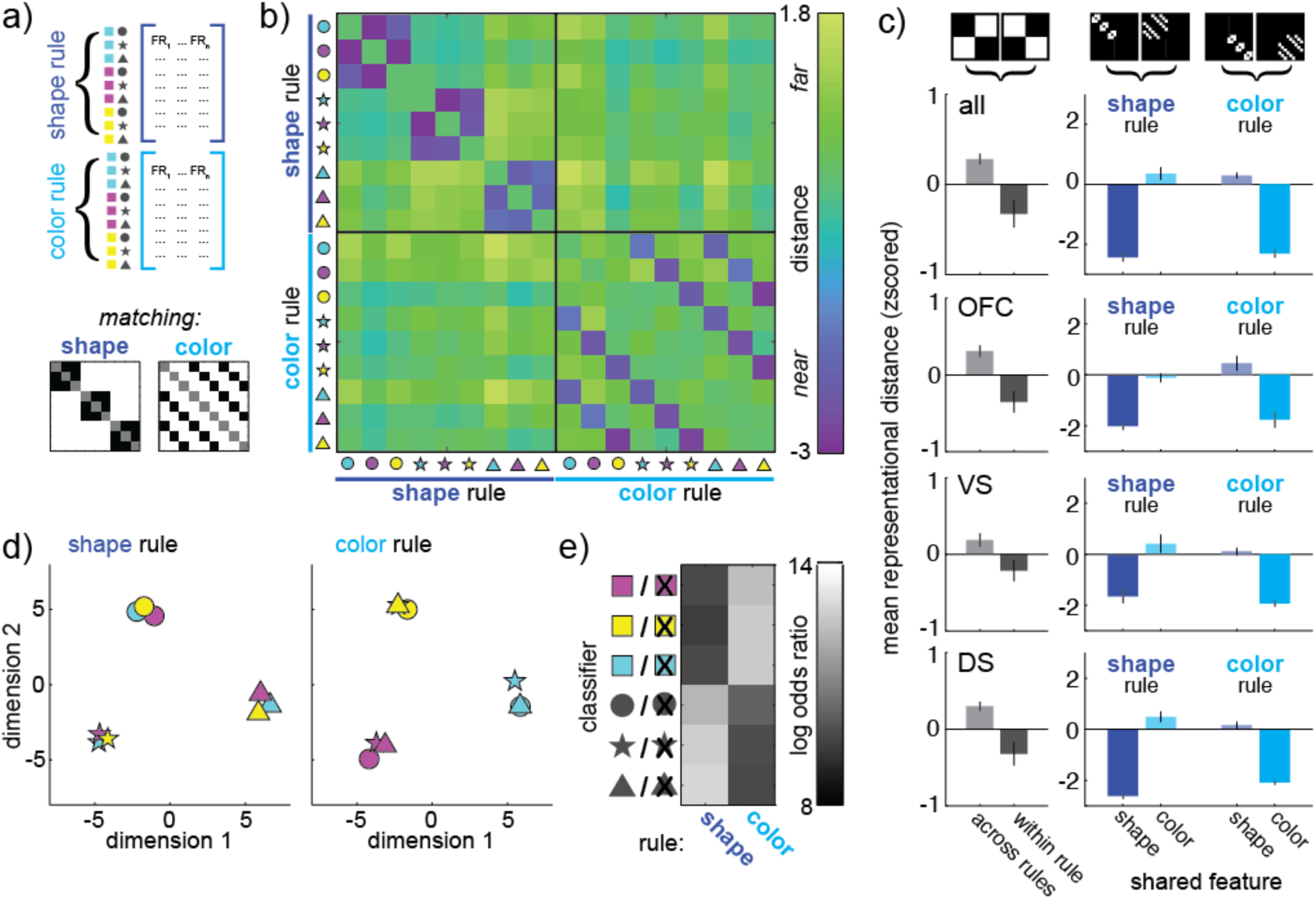
Changes in choice representations during color and shape rules. A) Top) To contrast shape-rule and color-rule choice representations, separate pseudopopulations are constructed for each class of rule. Bottom) The expected outcome if only shape (left) or color (right) influenced representational similarity. B) The representational similarity matrix for both shape-rule (upper left) and color-rule (bottom-right) decisions, all regions. C) Contrasts across the representational similarity matrix, calculated for all regions (top), then separately for OFC, VS, and DS (top to bottom). Left column) A change in representation between color-rule and shape-rule decisions would increase distance between representations measured across decision types (light gray), compared to within decision types (dark gray). Right column) Mean distance between stimuli that share the rule-relevant feature (saturated color) and rule-irrelevant feature (dim color), during shape-rule (left colum) and color-rule (right column) decisions. D) Multidimensional scaling plots illustrate how choice representations shift across shape-rule (left) and color-rule (right) decisions. Dimensions calculated separately for shape- and color-rule trials. E) The log odds ratio (chosen – unchosen) for each choice category classifier (y-axis) across shape and color rules (x-axis). Classifiers trained and fit to the pseudotrial matrix. Lighter shades = more confidence in the correct classification.

### Rule-relevant coding dimensions expand and rule-irrelevant coding dimensions compress during rule-based decision-making

There are multiple ways to increase the representational similarity of choices that differ only in rule-irrelevant dimensions. In the warping hypothesis, these results could occur because compressing irrelevant choice dimensions collapses any differences in the representation of choices that differ in these dimensions. However, an increase in similarity could also be due to an abstract rule identity signal. This is because simply encoding a blue-rule during blue star, circle, and triangle choices could make the representation of these choices more similar, regardless of shape. Thus, to meet the more stringent predictions of the warping hypothesis, rules must change the way that choices are represented along the dimensions of neural activity that are used to compute choice (see Methods)—not through a trivial effect of rule-encoding.

Subjects made the same choices for three different reasons which occurred with approximately equal frequencies. For example, they could make a blue-star choice because they were following a blue rule, following a star rule, or making rule-free decisions (**Figure 6A**). This meant that it was possible to identify the axes within neuron-dimensional space that predicted blue choices, regardless of the goal, by marginalizing across these goals. We used targeted dimensionality reduction to identify the directions in neuron-dimensional space that predicted choice category (**Figure 6B**; see Methods). This identified three coding dimensions that predicted choice color and three coding dimensions that predicted choice shape—where coding dimensions are the linear combinations of neuronal firing rates that best predicted changes in the log odds of one choice category. As expected, choice-predictive coding dimensions were neither identical nor orthogonal to the matching rule coding dimensions (mean angle between rule and matching choice coding dimensions = 52.7 degrees, range across choice categories = [48.5, 56.2]). This implies that rule-adherence did affect how choices were represented along choice-predictive coding dimensions, but that changes in choice-predictive coding dimensions could not be fully explained by encoding of the rule identity.

**Figure 6.**
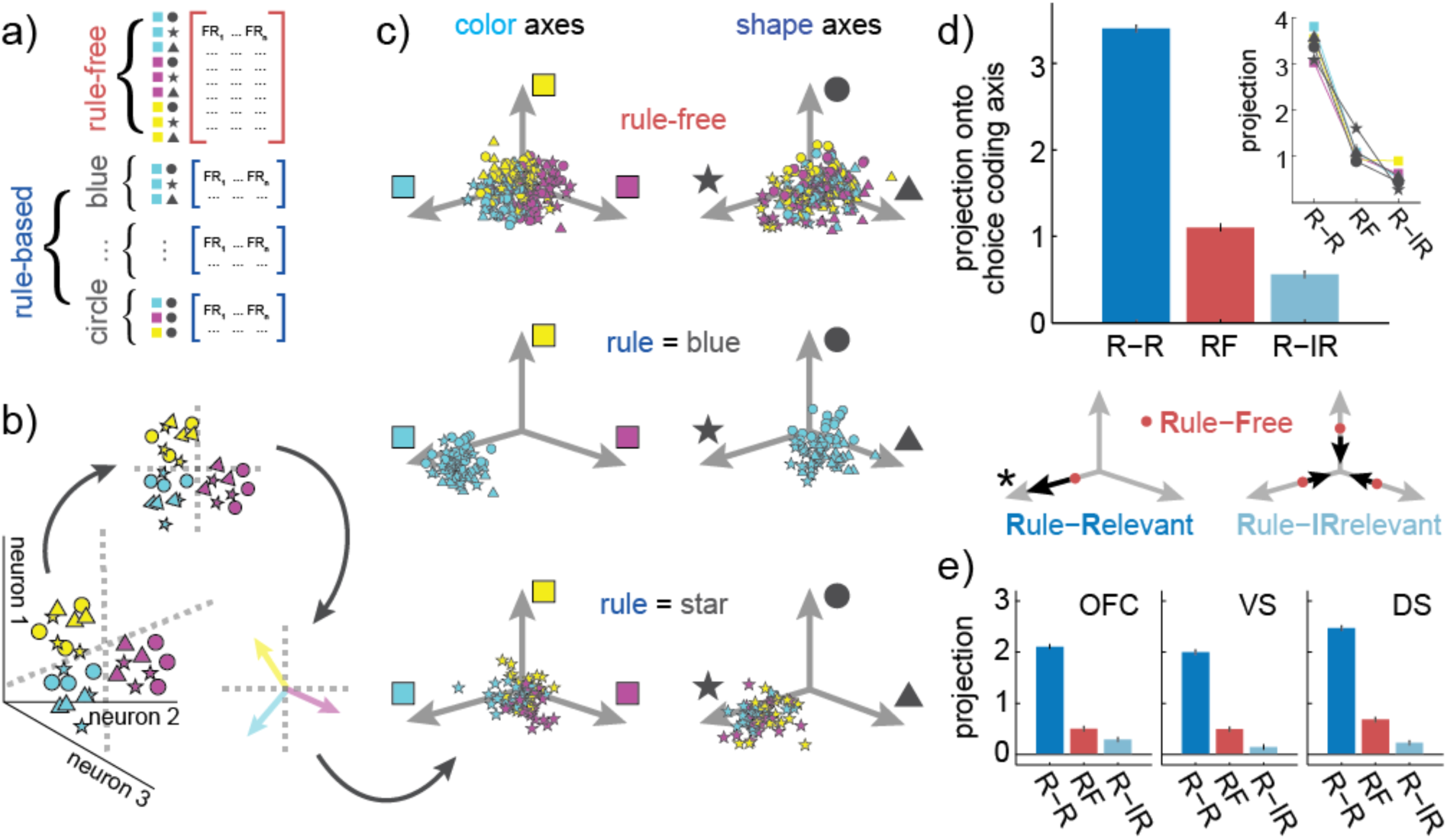
Changes in choice representations within a choice-predictive subspace. A) To determine how adhering to a specific rule changed choice representations within a choice-predictive subspace, pseudopopulations are constructed for each rule-based and rule-free state. A) Choice-predictive subspaces are identified with multiple logistic regression. This finds the separating hyperplanes (dotted gray lines) that best separate each class of choices in neuron-dimensional space. The cartoon illustrates two color hyperplanes, one separating magenta choices from other choices and one separating yellow choices from other choices. Pseudotrial activity is then projected into the subspaces defined by the color and shape hyperplanes. Within these subspaces, specific vectors correspond to the coding dimensions in neural activity that predict whether the choice will include each color or shape. These coding dimensions are included as reference vectors in the following panel. C) Distribution of rule free (top), color rule (middle), and circle rule (bottom) pseudotrials in each of the color (left column) and shape (right column) subspaces. D) The projection onto the coding dimensions of the chosen features for rule-free decisions (red, RF) and rule-based decisions (blue), with the latter separated according to whether the coding dimension is rule-relevant (dark blue, R-R, or rule-irrelevant (light blue, R-IR). Inset) Average projections for each color and shape rule, plotted separately. Bottom) A cartoon illustrating the central effect: compared to rule-free trials (red dots), rule-based decision-making pushes choice representations further along the rule-relevant axis (*), while compressing the rule-irrelevant subspace. E) Same as D, plotted separately for OFC, VS, and DS.

To understand how choice representations changed along choice-predictive coding dimensions, we projected each pseudotrial into the subspace defined by these coding dimensions. Formally, this meant that we found a low-dimensional projection of neural activity where position corresponded to the decoded probability of choice. For illustration, the three choice color coding dimensions were grouped into color subspace, while three choice shape coding dimensions were grouped into shape subspace (**Figure 6C**). Projections onto chosen-feature coding dimensions were substantially higher than projections onto unchosen feature coding dimensions (chosen mean = 1.69, range across choice categories = [1.55, 1.81]; unchosen mean = −1.54, range = [−1.64, −1.35]; p < 0.0001, t(1,3238) = 67.29, two-sample t-test, 95% CI = [3.13, 3.32]). Thus, the projection of each pseudotrial onto these choice-predictive axes did indeed predict choice.

However, the magnitude of the choice coding dimension projections differed, depending on the subjects’ goal: whether they chose the stimulus because the category was relevant to the rule, irrelevant to the rule, or the result of rule-free decision-making (**Figure 6D**). The representation of rule-relevant choice features was pushed furthest along the choice-predictive axis (mean = 3.40, range across categories = [3.03, 3.59]; sig. greater than rule-irrelevant: p < 0.0001, t(1,718) = 44.1, 95% CI = [2.72, 2.97], two-sample t-test; sig. greater than rule-free: p < 0.0001, t(1,718) = 35.1, 95% CI = [2.17, 2.43]). The next largest projection was the rule-free trials (mean = 1.10, range across categories = [0.89, 1.62]; sig. greater than rule-irrelevant: p < 0.0001, t(1,718) = 8.13, 95% CI = [0.41, 0.67], two-sample t-test). Finally, rule-irrelevant features had the smallest coding dimension projection, meaning that they were the least discriminable (mean = 0.56, range = [0.29, 0.90]). Thus, relative to how choices were represented during rule-free decisions, rule-based decision-making expanded the rule-relevant dimensions of choice representational space, but compressed the rule-irrelevant dimensions.

Next, we asked whether or not these same effects were apparent in all three regions individually. We found nearly identical results in each of OFC, VS, and DS (**Figure 6E**; see **Table S2 and S3**). Thus, together these results suggest that there was a distributed change in choice representations during rule-based decision-making such that choice dimensions that were consistent with the rule were expanded and irrelevant dimensions were compressed across a distributed network of regions linked to decision-making.

## DISCUSSION

We developed a new approach to defining rule-based and non-rule-based decisions in a dynamic rule-based decision task and found that rule-based decisions made more efficient use of limited neural resources than non-rule-based decisions. Fewer spikes were needed to generate rule-based choices across a distributed network of decision-making regions. Further, although we know that reducing neural activity tends to reduce information (Levy & Baxter, 1996), choice information was actually increased during rule-based choices. One way that implementing a rule could improve the efficiency of decision-making is by changing how decision-making problems are represented in the brain (Gigerenzer & Todd, 1999; Shah & Oppenheimer, 2008; Tversky, 1972). Indeed, we found distributed changes in neuronal representations that are broadly consistent with this hypothesis. In three brain regions, OFC, VS, and DS, adhering to a rule warped the geometry of neural decision-making space in two ways: it expanded neuronal choice representations along rule-relevant choice coding dimensions, while compressing representations along rule-irrelevant ones.

Representational changes during rule-based decision-making were remarkably similar across OFC, DS, and VS. Previous studies have focused on the differences between these regions (O’Doherty et al., 2004; Poldrack et al., 2001; Valentin et al., 2007), and, indeed, we did observe two striking differences between the regions here. First, we found that population activity in VS was more sparsely distributed across neurons during rule-based decision-making, compared to rule-free decision-making. Conversely, in OFC and DS, activity was sparser during rule-free decision-making. Because the BOLD signal, in particular, may be sensitive to the total number of activated neurons (Logothetis, 2008), it is possible that this dissociation could explain why studies using BOLD imaging might see an increase in BOLD signal during rule-based decisions in OFC and DS, but a decrease in BOLD in VS (O’Doherty et al., 2004; Poldrack et al., 2001; Valentin et al., 2007). We saw a decrease in spike counts in all three regions here, implying that hemodynamic responses may track sparseness measures more than they track average firing. Second, we found that the representational space of choices was more compact during rule-based decisions than rule-free decisions in OFC, but in VS and DS, adhering to a rule expanded the space of choice representations—increasing the distance between both similar and dissimilar choices in the striatum, but not the cortex.

One interpretation of these results is that stimulus-response rules are implemented, in part, through an attentional gate. This gate selects which features of choice options to prioritize for processing at the expense of others (Desimone and Duncan, 1995; Miller and Cohen, 2000; Pastor-Bernier and Cisek, 2011). Here, rules not only gate which stimuli are represented in the neural code (Rainer et al., 1998), but also *how* different stimuli are represented. This representational warping could be a powerful way to facilitate good decision-making by making it easier to classify options with respect to the attributes that matter to the decision-maker in the moment (cf. Machens et al., 2005). Of course, an attentional gate is not necessary for good decision-making and we are more than capable of representing every dimension of the decision-making problem (Mante et al., 2013). However, these results suggest that we do, under some circumstances, take advantage of a gating strategy that may reduce the energetic costs of decision-making.

In some cognitive theories, simple policies like rules are thought to reduce the computational costs of coming to a decision largely because they eliminate the need to either represent or to perform computations on rule-irrelevant information. However, here, the compression in rule-irrelevant coding dimensions was matched, if not exceeded, by expansion in rule-relevant dimensions. If there was some conserved decision-making quantity—like a fixed amount of evidence or value that must be accumulated to come to a choice—then the expansion of rule-relevant dimensions could be an inevitable consequence of compression in irrelevant dimensions. This is because eliminating the encoding of rule-irrelevant information would shift the choice process to over represent rule-relevant information in order to meet the fixed threshold for generating a decision. Of course, it is not clear that value-based decisions depend on an integrate-to-bound process (Gold & Shadlen, 2007; Rangel & Hare, 2010), and a bottleneck in visual attention could also lead to this type of conservation of evidence. Alternatively, it is also possible that rule-relevant coding dimensions are expanded because expansion facilitates critical rule-relevant computations, like the ability to classify the choice with respect to the current rule.

The idea that higher-order cognitive information can reshape sensory representations to facilitate specific computations has support in both influential theories of prefrontal function (Miller & Cohen, 2001) and modern views at the intersection of working memory and selective visual attention (Cukur et al., 2013; Myers et al., 2015; Myers et al., 2017). However, previous studies have not reported evidence that task rules can change the way that sensory features are represented in the brain. Future work is necessary to determine whether the critical difference between our study and these previous studies is one of task design, analysis method, or the fact that this study targeted limbic regions commonly implicated in decision-making, rather than prefrontal or extrastriate regions linked to controlling or executing visual attention (Katzner et al., 2009; Lauwereyns et al., 2001; Mante et al., 2013; Sasaki & Uka, 2009).

Certainly, one straightforward way to produce the kinds of broadly distributed representational changes we report here would be through a change in visual attention, either at the level of the extrastriate cortex (Martinez-Trujillo & Treue, 2004; Maunsell & Treue, 2006; Treue & Trujillo, 1999) or the prefrontal regions that direct featural attention (Bichot et al., 2015). Early featural selection could alter the way that stimulus features are represented within every region that receives ascending visual information (Ebitz and Moore, 2019). This would imply that mechanisms relevant for sensory and association systems may also apply to prefrontal regions and their striatal targets (Yoo and Hayden, 2018; Eisenreich et al., 2017). However, the effects we report here could also arise through other selective mechanisms, such as a bias coming directly into OFC, VS, and/or DS from dorsolateral prefrontal cortex (Miller & Cohen, 2001), an adjustment in gain from dorsal anterior cingulate (Ebitz & Hayden, 2016; Ebitz & Platt, 2015), or amplification through recurrent interactions between these regions and/or through the thalamus (Halassa & Kastner, 2017). Future work will be needed to determine whether the representational changes that occur during rule-based decisions are mediated by changes in selective visual attention or if gating only emerges at the level of these decision-making regions.

The brain is a highly evolved organ, and it should not be surprising if its processing is directed toward efficiency in complex and rapidly changing naturalistic tasks (Krakauer et al., 2017; Datta et al., 2019 Pezzulo and Cisek, 2016; Pearson et al., 2014; Calhoun and Hayden, 2014; Hayden, 2018). Rule adherence is likely to be an intrinsically costly operation, and would benefit especially from mechanisms that make it more efficient. While the CSST is not particularly naturalistic, it is somewhat more complex than standard laboratory decision-making tasks that involve a static rule set. As such, conventional tasks do not offer the opportunity to show neuronal effects of rule-based and rule-free decisions.

## METHODS

### Surgical procedures

All animal procedures were approved by the University Committee on Animal Resources at the University of Rochester and were designed and conducted in compliance with the Public Health Service’s Guide for the Care and Use of Animals. Two male rhesus macaques (*Macaca mulatta*) served as subjects. Standard surgical techniques, described previously (Strait et al., 2014), were used to implant a small prosthesis for holding the head and Cilux recording chambers (Crist Instruments). Chamber positions were verified by magnetic resonance imaging with the aid of a Brainsight system (Rogue Research). Neuroimaging was performed prior to surgery at the Rochester Center for Brain Imaging on a Siemens 3T MAGNETOM Trio Tim using 0.5 mm voxels. Animals received appropriate analgesics and antibiotics after all procedures and chambers were kept sterile with regular antibiotic washes and sealed with sterile caps. All data presented here were collected previously and have been used in earlier work (Sleezer et al., 2016, 2017; Sleezer & Hayden, 2016; Yoo, Sleezer, & Hayden, 2018). All analyses presented here are new.

### Electrophysiological techniques

OFC (115 neurons), VS (103 neurons), and DS (204 neurons) were approached through a standard recording grid (Crist Instruments) using a standard atlas to define OFC (Area 13M), VS (the core of the nucleus accumbens), and DS (both the caudate and medial part of the putamen). Specific coordinates each recording site have been described in detail previously (Sleezer et al., 2016). Recording locations were verified through a combination of Brainsight guidance and listening for characteristic sounds of white and gray matter while lowering the electrodes, which in all cases matched the boundaries predicted by the Brainsight system.

During each session, between 1 and 4 single electrodes (Frederick Haer; impedance range 0.8 – 4 M) were lowered using a microdrive (NAN Instruments) until the waveforms of single neuron(s) were isolated. Neurons were selected for study solely on the basis of the quality of isolation, not on task-related response properties. Cells were isolated and recorded with a Plexon system.

### General behavioral techniques

Animals were habituated to laboratory conditions and then trained to perform oculomotor tasks for liquid reward before training on the task. Previous training history for these subjects included two types of foraging tasks, intertemporal choice tasks, two types of gambling tasks, attentional tasks, and a basic form of a reward-based decision task. Eye position was sampled at 1000 Hz by an infrared eye-monitoring camera system (SR Research). Stimuli were controlled by a computer running MATLAB (The MathWorks) with Psychtoolbox and Eyelink Toolbox. Visual stimuli were presented on a computer monitor placed 57 cm from the animal and centered on its eyes. A standard solenoid valve controlled the duration of juice delivery. The relationship between solenoid open time and juice volume was established and confirmed before, during, and after recording.

These subjects had never been trained on tasks in which rule-reward mappings changed rapidly and required constant learning. Specifically, all previous training involved dynamic tasks in which risky or certain options varied unpredictably across trials or across short blocks. Subjects were, however, familiar with several tasks. These included a diet selection task (Blanchard & Hayden, 2014), a token gambling task (Azab & Hayden, 2017; Azab & Hayden, 2018; Farashahi et al, 2018), and several riskless choice tasks (Pirrone et al, 2018; Heilbronner and Hayden, 2016). In all of these tasks, there was no consistent relationship between spatial location and reward, and thus, subjects had no past opportunity to form a link between rewards and spatial positions.

### Behavioral task

This task has been described in detail previously (Ebitz et al., 2019; Sleezer et al., 2016; Sleezer & Hayden, 2016). This present task is a version of the CSST: an analogue of the WCST that was developed for use in nonhuman primates (Moore et al., 2005). Task stimuli are similar to those used in the human WCST, with two dimensions (color and shape) and six specific rules (three shapes: circle, star, and triangle; three colors: cyan, magenta, and yellow; **Figure 1A**). Choosing a stimulus that matches the currently rewarded rule (e.g. any blue shape when the rule is blue; any color of star when the rule is star) results in visual feedback indicating that the choice is correct (a green outline around the chosen stimulus) and, after a 500 ms delay, a juice reward. Choosing a stimulus that does not match the current rule results in visual feedback indicating that the choice is incorrect (a red outline around the chosen stimulus), and no reward for the trial.

The rewarded rule was fixed for block of trials. At the start of each block, the rewarded rule was drawn randomly. Blocks lasted until monkeys achieved 10, 15, 20, or 30 correct responses that matched the current rule (fixed across sessions). This meant that blocks lasted for a variable number of total trials, determined by both how long it took monkeys to discover the correct objective rule and how effectively monkeys exploited the correct rule, once discovered. Block changes were uncued, although reward-omission for a previously rewarded option provided unambiguous information that the reward contingencies had changed.

On each trial, three stimuli were presented asynchronously. One stimulus was presented at each of three locations on the screen. The color, shape, position, and order of stimuli were randomized. Stimuli were presented for 400 msec and were followed by a 600-msec blank period. (The blank period is omitted in **Figure 1A** because of space constraints). Monkeys were free to look at the stimuli as they appeared, which they typically did (Sleezer & Hayden, 2016). After the third stimulus presentation and blank period, all three stimuli reappeared simultaneously with an equidistant central fixation spot. When they were ready to make a decision, monkeys fixated on the central spot for 100 msec and then indicate their choice by shifting gaze to one stimulus and maintaining fixation on it for 250 msec. If the monkeys broke fixation within 250 milliseconds, they could either again fixate the same option or change their mind and choose a different option, although they seldom did so. Thus, the task allowed the monkeys ample time to deliberate over their options, come to a choice, and even change their mind, without penalty of error.

### General data analysis techniques

Data were analyzed with MATLAB. Two sample t-tests were used for all pair-wise statistical comparisons except for the mutual information analyses because these samples visibly deviated from the normality assumptions of the test. Neural activity was analyzed in the fixed 3350 ms epoch that started at the presentation of the first option and ended just before feedback was provided. Firing rates were z-scored within neurons for all analyses, except in direct comparisons of total firing rate across rule-based and rule-free decisions (e.g. **Figure 2**), where firing rates were centered, but not scaled. All pseudopopulation results are reported for a single, randomly-seeded pseudopopulation, but were confirmed with 1) different random seeds, and 2) bootstrap tests across 1000 random seeds.

### Identifying information-maximizing choices

We determined how much information would be gained from different choice strategies with a model that used the recent history of rewards and choices to estimate the likelihood that each feature was the correct one (Ebitz et al., 2019). The influence of past outcomes and choices decays exponentially (Lau & Glimcher, 2005), so we took advantage of the fact that the last trial has the single largest influence on choice to construct a simplified model with a 1-trial history. Assuming all possible histories of choices and rewards before the last trial (X_1:t-2_), we initialize a uniform prior that each feature (f) of the N_f_ features is the correct feature (f*):

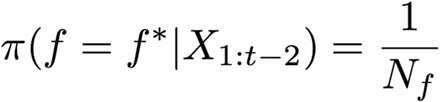

After a choice is made at time t-1, we calculate the likelihood that the chosen feature was correct in a reward-dependent fashion. If the choice is rewarded (r=1), the likelihood is:

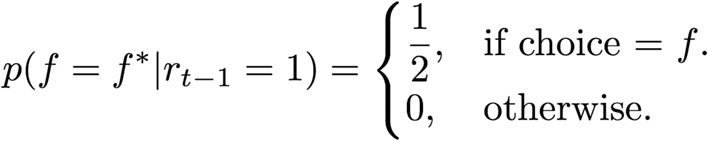

If the choice is not rewarded, the likelihood is:

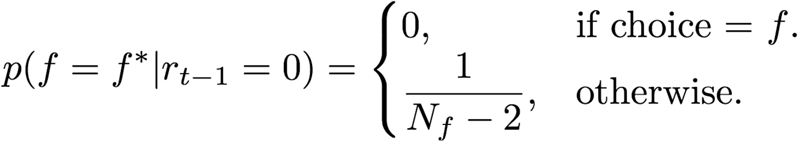

Multiplying the prior and likelihood and renormalizing into a valid probability distribution would give the posterior probability that each feature is best after t-1, which is also the new prior at the start of the rule-free trial t. Because we knew, on average, whether or not the monkeys had been rewarded for the last choice before a rule-free decision, we calculate the average prior for rule free decisions by taking an average of these two likelihoods, weighted by the probability that they occurred:

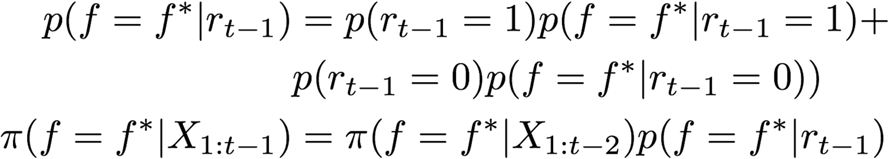

To determine how different rule free choices would change the information about what choice is best, we first calculate the entropy of the prior:

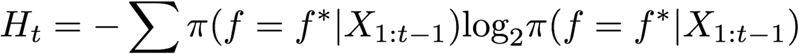

Then, we calculate the posterior distribution that we would observe after rule-free choices. We first calculate the likelihoods, as described above, for all possible combinations of choice (c_t_) and reward outcome (r_t_), then multiply each likelihood with the prior to get all possible posterior distributions after the rule-free choice:

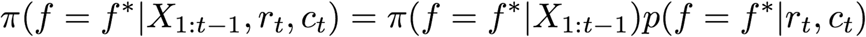

There were two possible outcomes of each choice—one where the animal would be rewarded, and one where they would not—and the likelihood of these futures differed systematically across the different choices. Therefore, the estimated posterior entropy for each choice is an average of these possible futures, weighted by their likelihood of occurring:

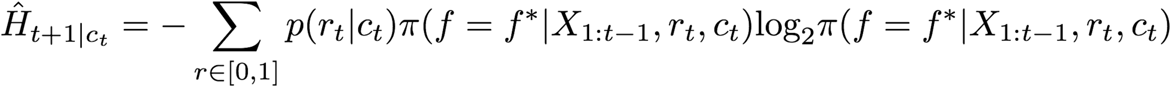

Where the probability of reward for each choice is just the probability that the choice will include the best feature:

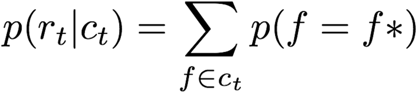

The information maximizing choice is then one that would cause the largest drop in the model’s uncertainty about what feature is correct. That is, it would be the choice, c, that maximizes:

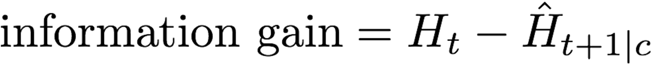

Small amounts of noise, |N(0, 10^−4^)|, were added to all 0’s so that information would be computable. Choices were then classified as differing in 0, 1, or 2 stimulus features from the last choice and the average information gain across these classes is illustrated in the inset of **Figure 1G**.

### Hidden Markov model (HMM)

A hidden Markov model was used to infer the latent goal state underlying each choice. This model allowed us ask whether, for example, each blue-star choice was made because 1) it was a star, 2) it was blue, or 3) the monkey was searching for the correct feature. We have previously used this method to identify latent goal states in other tasks (Ebitz, Albarran, & Moore, 2018) and described how this model was developed for this task (Ebitz et al., 2019).

Briefly, in an HMM framework, choices (y) are “emissions” that are generated by an unobserved decision process that is in a latent, hidden state (z). Each hidden state has some observation model, which dictates the probability of choosing each option when the process that state. For example, the blue-rule state had an observation model where the probability of choosing the blue option was 1, but the probability of choosing a non-blue option was 0. Formally:

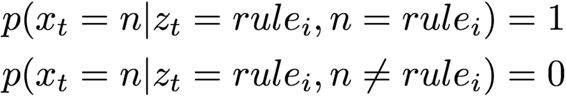

Conversely, because we do not know what monkeys would choose in a rule-free state, the observation model for any of the N=3 choice options (*n)* during rule-free states is:

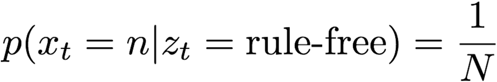

In addition to these observation models, the hidden states are also defined by how long the process will tend to stay in that state and where the process will tend to go from that state. Because the model is Markovian (the current state is conditionally independent of all previous history, given the last state), these properties are described by a stochastic transmission matrix, whose entries are the probability of transitioning from each state to every other state, including itself. Monkeys could not divine the new rule following a change point and instead had to explore to discover it, so direct transitions between different rule states were fixed at 0. For the same reason, the monkeys were also assumed to start in the rule-free state on the first trial each day. Transmission parameters were tied across the six rule states, meaning that the model ultimately had only 2 free parameters: the probability of staying in the rule-free state and the probability of staying in any of the six rule-based states.

These two parameters were fit to each individual session via expectation-maximization using the Baum Welch algorithm (Bilmes, 1998; Murphy, 2012). This algorithm finds a (possibly local) maxima of the complete-data likelihood, which is related to the joint probability of the hidden state sequence Z and the sequence of observed choices Y. The algorithm was reinitialized with random seeds 100 times, and the model that maximized the observed (incomplete) data log likelihood was ultimately taken as the best for each session. We then used the Viterbi algorithm to identify the most probable a posteriori latent state underlying each choice (Murphy, 2012), given the sequence of choices and the model we fit.

### Mutual information

To estimate the amount of choice information in single neurons, we calculated mutual information between firing rate (r) and choice identity (c) as:

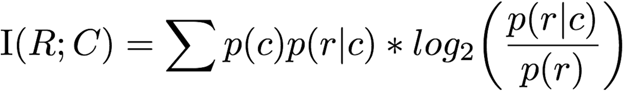

Firing rates were discretized via quantile binning within neurons to allow us to calculate these probabilities directly. Two firing rate bins per neuron were used because maximized the number of observations per condition (combination of stimulus identity and firing rate), though the results were similar with more firing rate bins.

The monkeys completed fewer rule-free trials than rule-based ones, and mutual information estimates are systematically inflated when the number of observations is low (Panzeri et al., 2007) due to limited-sampling biases. We controlled for this bias in two ways. First, we randomly downsampled the rule-based trials to match the count of rule-free trials and report the results of this downsampled distribution in the text and figure. Second, we calculated the expected mutual information under the hypothesis of no relationship between choice and firing rate in each case, including these as reference lines in the plot.

### Sparsity and the Gini index

The Gini index is a measure of the sparsity of the distribution of some resource (in this case, spikes) across a population (in this case, neurons). Here, a sparse distribution is one in which a disproportionate number the total population spikes were emitted by a small number of neurons. The Gini index has a few nice properties for measuring spike sparsity in that it is scale-invariant in terms of both the number of spikes and the number of neurons and it is sensitive to the addition of long-tail outliers, both in terms of very low and very high firing neurons (Hurley & Rickard, 2009).

Given a sorted vector of mean firing rates for n neurons (a sorted pseudotrial, see below), x = [x_1_, x_2_ … x_n_], where x_1_ ≤ x_2_ ≤ x_n_, the Gini index is calculated as:

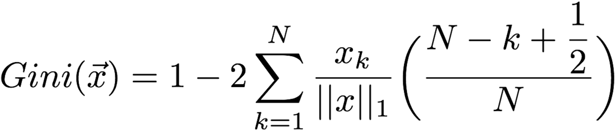

To give an intuition for this measure, the Gini index quantifies the area under the curve in a plot of the cumulative total population spikes against the cumulative total number of neurons (**Figure S1A**). If each neuron contributed equally to the total spike count, this plot would follow unity and the Gini index would equal 0.5. As more and more neurons are silent (equivalently, as more of the spikes are concentrated in a smaller number of neurons), this curve will deviate more from the unity line and the Gini index will approach 1. The Gini index was calculated for each pseudotrial, using the same pseudotrial matrix illustrated in Figure 4.

### Pseudopopulation choice representations

To estimate how choice representations at the population level changed with rules, we combined all the neurons into pseudopopulations. That is, we treated separately recorded neurons as if they were simultaneously recorded (Churchland et al., 2012; Machens et al., 2010; Mante et al., 2013; Meyers et al., 2008). Though this pseudopopulation approach does not allow us to reconstruct the covariance structure between simultaneously recorded neurons, it can still be useful for generating first order insights into how population activity changes across various conditions.

Within each task condition (combination of chosen color, shape, and state), firing rates from separately recorded neurons were randomly drawn with replacement to create a pseudotrial firing rate vector for that task condition, with each entry corresponding to the activity of one neuron in that condition. Together, these pseudotrial vectors were stacked into the trials-by-neurons pseudopopulation matrix. Twenty pseudotrials were drawn for each condition, based on the observation that approximately 75% of conditions had at least this number of observations (mean per neuron per condition = 28, median = 26). If a small number of conditions were missing for a particular neuron (<5), we imputed the neurons’ mean firing rate. If a large number of conditions were missing, the neuron was excluded from these analyses. Neurons were also excluded if their mean firing rate was less than 2 spikes/second (Mante et al., 2013). Four of 422 neurons were excluded by these criteria, all of which were in VS. All reported pseudopopulation results come from a single pseudopopulation, but were confirmed by bootstrap tests across 1000 randomly re-seeded pseudopopulations.

We constructed a total of three different pseudotrial matrices, as illustrated in Figures 5, 6, and 7. These differed only in how we defined “state”. The first pseudopopulation (**Figure 4**) was constructed without respect to the specific rule that monkeys were using (i.e. all rule-based blue-star choices were combined, regardless of whether the rule was choose-blue or choose-star). We took this approach because we had no knowledge of which, if any, stimulus features were relevant during rule-free decisions. By creating a rule-based pseudopopulation without respect to the specific rule, we equated our knowledge about the two conditions and isolated any changes in choice representations during rule-based and rule-free decision-making. The second pseudopopulation (**Figure 5**) included information about whether the rule was color-based or shape-based, but not any information about whether it was a circle, star, or triangle shape-based rule, for example. This allowed us to isolate the effects of rule domain on stimulus feature representations. Finally, the third pseudopopulation (**Figure 6**), included information about the specific rule that the monkeys were choosing (blue-rule, or circle-rule). This allowed us to determine how specific rules affected rule-relevant and rule-irrelevant choice dimensions.

### Representational Similarity Analysis

To measure the representational similarity between different choices in a pseudopopulation, we first calculated the geometric mean choice representation within combinations of choice identity and choice type. Then, we measured the Euclidian distance between all pairs of mean choice representations. This created a matrix of pairwise distances between choice representations (e.g. **Figure 4 and 5**) in arbitrary units, which was zscored for all reported analyses. All analyses and zscoring excluded the distance between the chosen stimulus identity and itself, which was 0 by definition. Representational similarity matrixes are illustrated as the average of 500 randomly seeded pseudopopulations, but all statistical tests were performed on a single pseudopopulation.

### Category and identity tuning in single neurons

To determine whether choice category (versus identity) tuning was increased in single neurons during rule-based decision-making, we first quantified the extend of stimulus identity tuning in single neurons with an ANOVA. Here, we modeled the firing rate *Y* on a given trial *k* where choice *i* was made as:

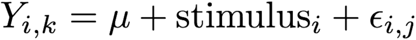

Where stimulus_i_ was identity-coded, meaning that each of the 9 unique possible combinations of color and shape were modeled independently. For each neuron, we then calculated the extent to which this model captured the variance in the neural data by calculating the F-test statistic of this ANOVA:

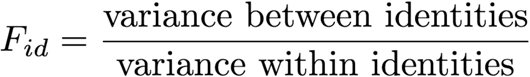

Next, we did the same with a model that independently modeled the contribution of the chosen stimulus color and shape to firing rate:

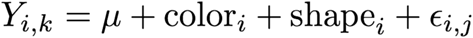

In this model, color and shape effects are assumed to be independent and additive. That is, this model assumes that there is no encoding of stimulus identity beyond what can be explained by separately knowing how a neuron responds to chosen color and shape. Again, we calculated how well this model captured firing rates via its F-test statistic:

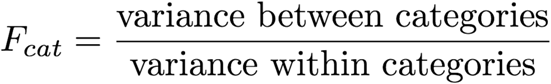

These two models were fit separately to each neuron during rule-based and rule-free decisions, respectively. We then calculated the relative extent of stimulus identity and stimulus category tuning during the two types of decisions with a “identity-category index”, which is a ratio of the F-test statistics of the choice identity and choice category models:

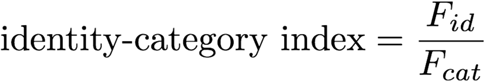

Here, a value greater than 1 would indicate that independently modeling each choices’ identity improved model fit, compared to modeling only choice category. Conversely, a value equal to or less than 1 would indicate that choice category information was sufficient to explain the variance in firing rate during decision-making.

### Choice predictive subspace analyses

To identify choice-predictive subspaces, we used a form of targeted dimensionality reduction based on multinomial logistic regression (Ebitz et al., 2018). Targeted dimensionality reduction is a class of methods for re-representing high-dimensional neural activity in a small number of dimensions that correspond to variables of interest in the data (Cohen & Maunsell, 2010; Cunningham & Byron, 2014; Mante et al., 2013; Peixoto et al., 2018). Thus, unlike principle component analysis—which reduces the dimensionality of neural activity by projecting it onto the axes that capture the most variability in the data—targeted dimensionality reduction reduces dimensionality projecting activity onto axes that encode task information or predict behavior.

Here, we were interested in how rule-based decision-making changed how choices were represented, so we first identified the axes in neural activity that predicted choice. The design of this task allowed us to dissociate choice-predictive axes from axes that encoded rule information because the same choice could be generated in three ways: via color-rule, shape-rule, or rule-free computations. Because these occurred in roughly equal proportion (33% of trials were rule-free, while 67% were distributed evenly across shape and color rules), we could identify the axes that predicted choice, regardless of why this choice was made.

We used multinomial logistic regression to find the separating hyperplanes in neuron-dimensional space that best separated choices to one category (i.e. blue) from other choices (i.e. not blue). Formally, we fit a system of six logistic classifiers:

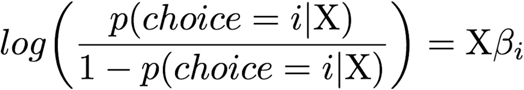

Where X is the trials by neurons pseudopopulation matrix of firing rates and β_i_ is the vector of coefficients that best differentiated neural activity on trials in which a choice matching category i is chosen from activity on other trials. The separating hyperplane for each choice i is the vector (a) that satisfies:

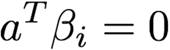

Meaning that β_i_ describes a vector orthogonal to the separating hyperplane in neuron-dimensional space, along which position is proportional to the log odds of choice. By projecting a pseudotrial vector x onto a coefficient vector β_i_, we are re-representing that trial in terms of its distance from the separating hyperplane corresponding to choice category i. Projecting that trial onto all six classifiers, then re-represents that high-dimensional pseudotrial in six dimensions— each one corresponding to the likelihood that the choice will include that feature as predicted by the population activity. Because only decisions within a feature domain were mutually exclusive, the logistic classifiers naturally grouped into two sets: the color-category classifiers and the shape-category classifiers. These defined two subspaces in the neural activity: one in which trial position predicted choice color and one where it predicted choice shape (**Figure 6**).

Separating hyperplanes were fit via regularized maximum likelihood (ridge regression). Regularization helps reduce overfitting by penalizing models with large coefficients. The extent of this penalty—the regularization parameter λ—was set to the minimum value that maximized cross validated classification accuracy (20-fold cross validation, training on pseudotrial matrices constructed from 90% of the trials and tested on matrices constructed from the 10% of trials that were held out; cross-validated accuracy evaluated at 25 log-spaced values for λ, range: [0, 10^4^]). Nearly identical results were observed across a wide range of λ values, including 0. Rule-encoding vectors were identified through the same procedure as choice-encoding vectors, but the classifiers were trained to predict the color or shape of the rule, rather than the color or shape of the choice.

---

**Supplemental Figure 1:**
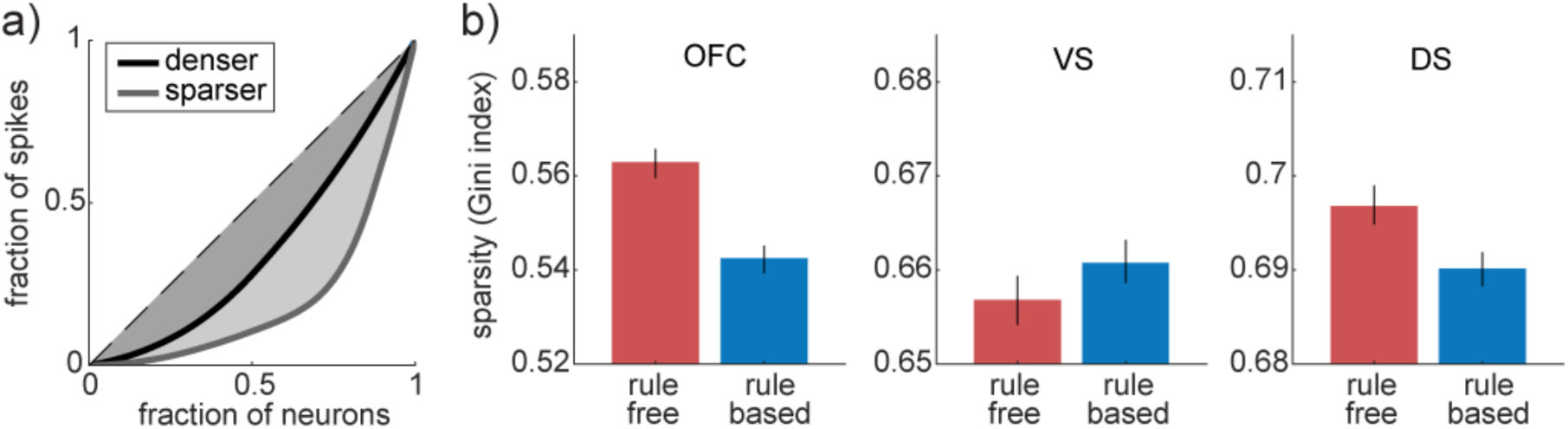
Heterogenous changes in the sparsity of neural activity across regions. A) The Gini index is a measure of the sparsity of the distribution of spikes across a neuronal population. As more spikes are concentrated in a smaller number of neurons (light trace), the area under the curve (shaded region) increases, increasing the Gini index. B) Sparsity changes between rule-free (red) and rule-based (blue) decision-making in VS (left), DS (middle) and OFC (right). Bars = STE across pseudotrials.

**Supplemental Table 1:**
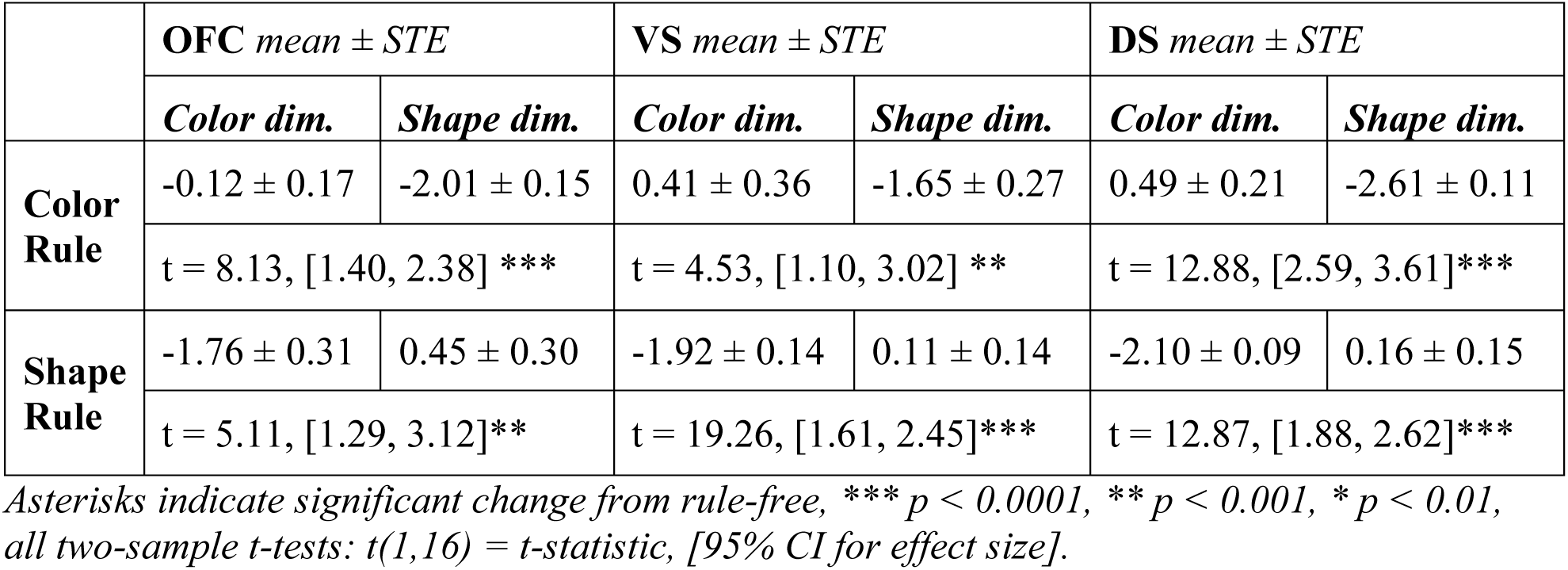

| | OFC mean ± STE | | VS mean ± STE | | DS mean ± STE | |
| --- | --- | --- | --- | --- | --- | --- |
|  | Color dim. | Shape dim. | Color dim. | Shape dim. | Color dim. | Shape dim. |
| Color Rule | -0.12 ± 0.17 | -2.01 ± 0.15 | 0.41 ± 0.36 | -1.65 ± 0.27 | 0.49 ± 0.21 | -2.61 ± 0.11 |
|  | t = 8.13, [1.40, 2.38] *** |  | t = 4.53, [1.10, 3.02] ** |  | t = 12.88, [2.59, 3.61] *** |  |
| Shape Rule | -1.76 ± 0.31 | 0.45 ± 0.30 | -1.92 ± 0.14 | 0.11 ± 0.14 | -2.10 ± 0.09 | 0.16 ± 0.15 |
|  | t = 5.11, [1.29, 3.12] ** |  | t = 19.26, [1.61, 2.45] *** |  | t = 12.87, [1.88, 2.62] *** |  |

Asterisks indicate significant change from rule-free, \*\*\* $p < 0.0001$ , \*\* $p < 0.001$ , \* $p < 0.01$ , all two-sample $t$ -tests: $t(1,16)$ = $t$ -statistic, [95% CI for effect size].

**Supplemental Table 2:**
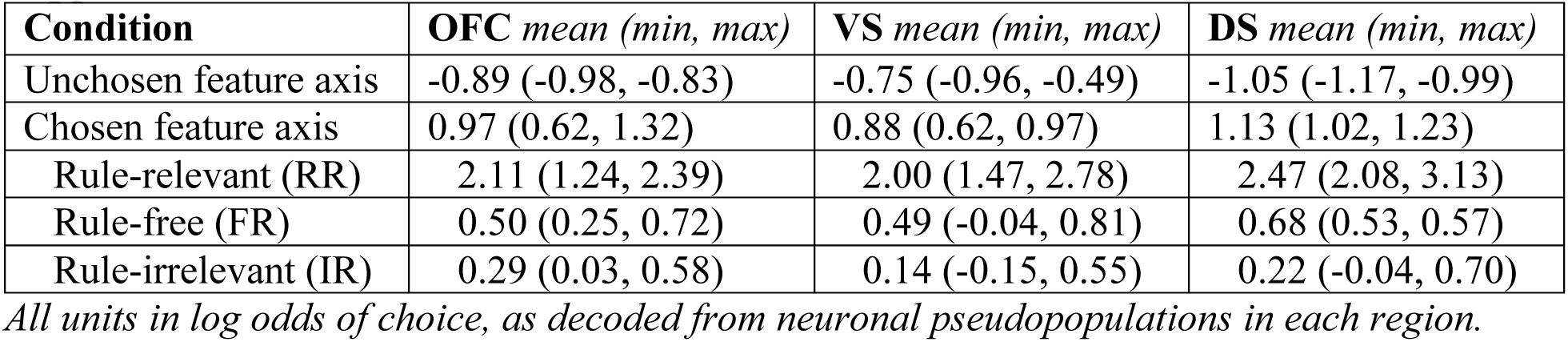

**Supplemental Table 3:**

| Test | OFC *t*-stat, 95% CI | VS *t*-stat, 95% CI | DS *t*-stat, 95% CI |
| --- | --- | --- | --- |
| Chosen ≠ Unchosen | *t* = 44.8, [1.78, 1.94]*** | *t* = 36.0, [1.53, 1.71]*** | *t* = 48.1, [2.09, 2.27]*** |
| RR ≠ FR | *t* = 21.8, [1.46, 1.75] *** | *t* = 20.0, [1.36, 1.65] *** | *t* = 23.4, [1.64, 1.94] *** |
| RR ≠ IR | *t* = 25.6, [1.68, 1.95] *** | *t* = 24.0, [1.70, 2.00] *** | *t* = 31.0, [2.10, 2.38] *** |
| IR ≠ FR | *t* = 2.9, [0.07, 0.35] ** | *t* = 4.49, [0.20, 0.50] *** | *t* = 6.2, [0.31, 0.60] *** |

*Asterisks indicate significant change from rule-free, \*\*\* $p < 0.0001$ , \*\* $p < 0.01$ , all two-sample *t*-tests: $t(1,718)$ = *t*-statistic, [95% CI for effect size], except chosen/unchosen, where $t(1,3238)$ .*

